# An open-source system for recording the auditory brainstem response (ABR) and an example application in mice with hearing loss

**DOI:** 10.64898/2025.12.17.694771

**Authors:** Rowan Gargiullo, Cedric Bowe, Abigail F McElroy, Jessica Mai, William N Goolsby, Chris C Rodgers

## Abstract

The auditory brainstem response (ABR) is a measure of the neurophysiological response to sound, widely used in clinical and research settings. In this manuscript, we describe an open-source and low-cost system suitable for measuring the ABR in mice. (This system is not evaluated or approved for clinical use in humans.) The heart of the system is the commercially available Texas Instruments ADS1299 chip, which is designed for multi-channel differential recording of biosignals. We designed auxiliary hardware and software to record and visualize the ABR. To demonstrate that this system is capable of high-quality and low-noise measurements, we recorded the response to free-field auditory clicks from the left and right. Next, we compared different electrode configurations, quantified the most consistent aspects of the ABR, and assessed variability within and across mice. Finally, we demonstrated how to detect the decreased auditory sensitivity caused by surgically induced conductive hearing loss. Users can deploy this open-source system at low cost and customize it for different applications, such as recording other biosignals like electromyography (EMG) or electrocardiography (ECG).

## Introduction

Neural circuits in the auditory system detect, localize, and identify sound, giving rise to the sense of hearing. The auditory brainstem response^1^ (***ABR***) is the sound-evoked bioelectric potential generated by electrophysiological activity in auditory brain regions. While intracranial recordings permit higher resolution, the ABR can be measured minimally- or non-invasively with extracranial electrodes placed near the ear. Measuring the ABR requires careful setup and signal processing because it is a small signal (<10 µV at the skin) embedded in a noisy background of other signals from non-auditory brain regions, the heart, the nearby muscles of the head and neck, and non-biological electrical noise from the environment.

The ABR has clinical and research applications. In clinical settings, it is used for routine assessment of auditory function in newborns and to diagnose certain forms of acquired hearing loss in children and adults^1^. In research settings, the ABR can be used to assess auditory function in humans^2^, laboratory mice^3–12^, rats^13–15^, chinchillas^16^, guineas pigs^16^, gerbils^16,17^, budgerigars^18^, naked mole-rats^19^, cats^20–22^, and cephalopods^23^. Moreover, the ABR is altered in some mouse models of Alzheimer’s disease^24–26^, suggesting that this signal may have diagnostic utility beyond hearing loss.

Many systems for measuring the ABR are proprietary or expensive. Open-source hardware can provide transparency, customizability, and lower cost, thereby broadening access to science^27–31^. There are also educational benefits to open-source hardware as users teach themselves how to assemble, use, and even improve the design. For these reasons, we set out to develop and test an open-source system for measuring the ABR built around the Texas Instruments ADS1299 (https://www.ti.com/product/ADS1299), a multi-channel differential amplifier chip designed for measuring biological signals. A few studies have described using the ADS1299 to measure the ABR in humans^32–34^ but generally without sharing complete hardware and software designs (although see ref^35^ for a well-documented example of multi-modal monitoring beyond ABR). Moreover, we are not aware of any studies using the ADS1299 to record ABR in laboratory animals or to detect hearing loss.

The present manuscript describes our open-source design and demonstrates how it can be used for high-quality recordings in mice. We provide a statistical description of the recorded ABR and other biosignals like the electrocardiogram (ECG), describe new metrics for visualizing and quantifying the ABR, and compare the results of different electrode locations. Finally, we illustrate how this system can be used to assess hearing loss. We have shared all of the data presented here on Zenodo^36^, and all of our software and hardware designs are freely available at https://github.com/Rodgers-PAC-Lab/ABR2025.

## System design

### Problem specification

The ABR arises from neural activity distributed over multiple brain regions. Specialized hair cells in the inner ear transduce incoming acoustic energy into neurophysiological responses in the afferent auditory nerve, which innervates the cochlear nucleus. From that brainstem nucleus, the ascending auditory pathway continues through the superior olivary complex, the lateral lemniscus, the inferior colliculus, the medial geniculate body in the thalamus, and the auditory cortex^37^.

The ABR is typically recorded on the skin close to each ear (near the auditory nerve) and at the cranial vertex (near the inferior colliculus; ***Fig 1a***). In this study, we used the abbreviations ***L*** and ***R*** to designate the points near left or right ear and ***V*** to designate the vertex. ***V*** is a useful point of comparison because its location on the midline means that any auditory-evoked activity would be expected to be the same for sound from the left or right, at least under the assumption that auditory processing is symmetrical.

**Figure 1.**
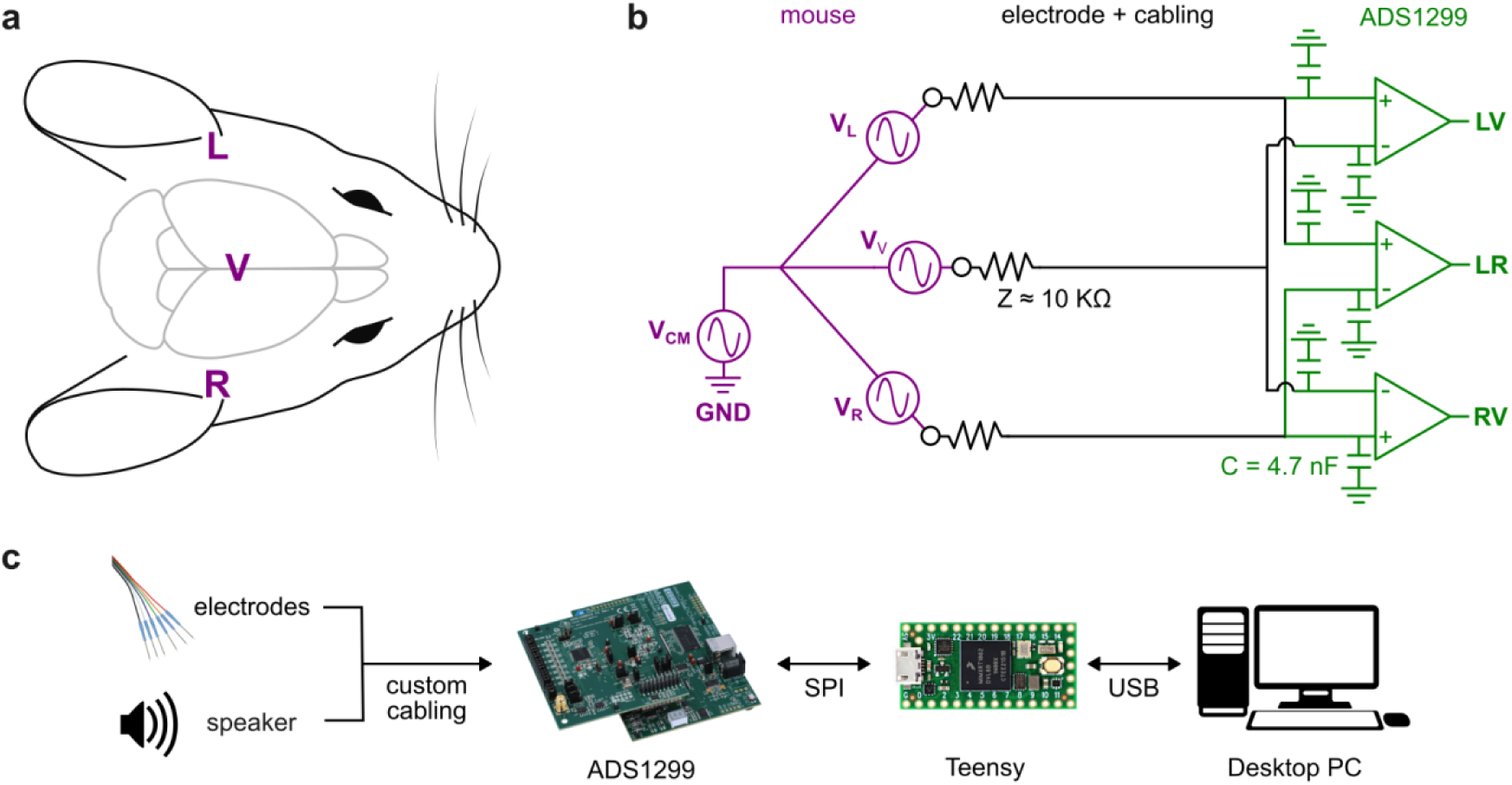
Schematic of overall approach. ***a)*** Approximate location of ***L***, ***V***, and ***R***. Image adapted from one by Luigi Petrucco^68^. ***b)*** *Purple*: ABR modeled as three voltage sources ***V_L_***, ***V_R_***, and ***V_V_*** in series with common-mode noise ***V_CM_***. *Black*: electrodes and cabling. *Green:* differential recording of ***LV***, ***RV***, and ***LR*** signals in the ADS1299. ***c)*** Recorded signals flow to ADS1299 to Teensy to desktop PC. Control signals flow the other way. Image credits: FreeSVG.org (speaker), LifeSync (electrodes), Texas Instruments (ADS1299), PJRC.com (Teensy), and VectorPortal.com (Desktop PC).

Schematizing this problem in an electrical circuit model, we treat the ABR as voltage sources at the left (***V_L_***) and right (***V_R_***) ears and a third voltage source at the vertex (***V_V_***; ***Fig 1b***). Although the mapping between neural activation and extracranial voltage is complex^38^, these model voltage sources represent the combined activation of multiple auditory-responsive brain structures. A common-mode noise source (***V_CM_***) arises from power line noise and other unwanted signals that are common between all three points. Generally, ***V_CM_*** is much larger than the ABR signals, which makes it difficult to record them.

### Differential recording

Differential recording is the standard approach to recording ABR or other biosignals with small-amplitude signals and large common-mode noise^39^. In this case, we placed subdermal needle electrodes at points ***L***, ***R***, and ***V***; a fourth electrode at the base of the tail was connected to electrical ground. With single-ended recording, we would record the signal at each point (e.g., ***L***) with respect to electrical ground. In contrast, with differential recording we record the voltage *difference* between two points (e.g., ***L*** and ***R***) by connecting one electrode to the positive input and the other to a negative input of a differential amplifier. The differential amplifier subtracts the negative input from the positive, which rejects (cancels out) common mode noise.

We recorded three differential signals, which we called ***LR*** (left minus right), ***LV*** (left minus vertex), and ***RV*** (right minus vertex). This configuration is redundant because any of the three signals can be computed from the other two. Previous studies often recorded only one of these differential signals at a time (e.g., ***LV***), but recording all three allowed us to compare their utility for testing auditory function. Although ***LR*** may be especially useful for measuring lateralized responses^6–8^, it is not as widely used as the vertex-ear configurations.

By convention, many (but not all^1^) laboratories use the vertex ***V*** as the positive (non-inverting) electrode and the ear as the negative (inverting) electrode. Because we were most interested in the signal at the ear, we used the opposite convention: whenever we recorded from ***V*** it was always with the negative electrode. This choice means that our reported signals will appear inverted from previous work using the opposite convention.

### System overview

We used the ADS1299 to perform differential amplification and digitization, a Teensy 4.0 microcontroller to send control signals and to receive and buffer data, and a standard desktop PC to save the data (***Fig 1c***). The ADS1299 provides a 16 kHz sampling rate and 8 differential inputs. Unlike other systems, this chip does not rely on high-gain analog amplification to deliver an acceptable signal-to-noise ratio, but rather on fine voltage resolution and low noise. With 24-bit analog-to-digital converters operating at the maximum gain of 24, the full range input is ±187 mV, the voltage resolution is 22 nV, the input referred noise is 1.66 μV rms, the common-mode rejection ratio is 120 dB, and the crosstalk between channels is -110 dB. (Explanation of these standard specifications is available in ref^39^.) Because biosignals vary greatly in amplitude (e.g., EMG is about 4 orders of magnitude larger than ABR), the important point for this manuscript is that these specifications allow recording over almost 7 orders of magnitude without clipping (187 mV / 22 nV ≈ 10^7^), although filtering or averaging is required for signals smaller than the input referred noise.

We designed a physical enclosure to house the ADS1299 and all auxiliary electronics, with banana plug breakout connections for differential inputs from the electrodes (***Supplemental Figure 1-4, Supplemental Table 1***). We used the ADS1299EEG-FE development kit (https://www.ti.com/tool/ADS1299EEGFE-PDK), comprising a MMB0 motherboard and an ADS1299EEG-FE daughterboard, but we used the motherboard only to provide power. The daughterboard included a passive analog pre-filter on each input channel: a 4.7 nF capacitor to ground, resulting in a lowpass filter of about 3.4 kHz assuming an electrode impedance of 10 kΩ (***Fig 1b***). The enclosure included a Teensy 4.0 microcontroller, which we programmed to send commands to and receive data from the ADS1299 over a 4-wire SPI protocol. For alignment with the audio stimulus, we used an additional differential input to record the voltage on the speaker. Alternatively, the Teensy could be used to collect digital pulses time-locked to stimulus onset. All required materials are listed in ***Table 1***.

**Table 1.**
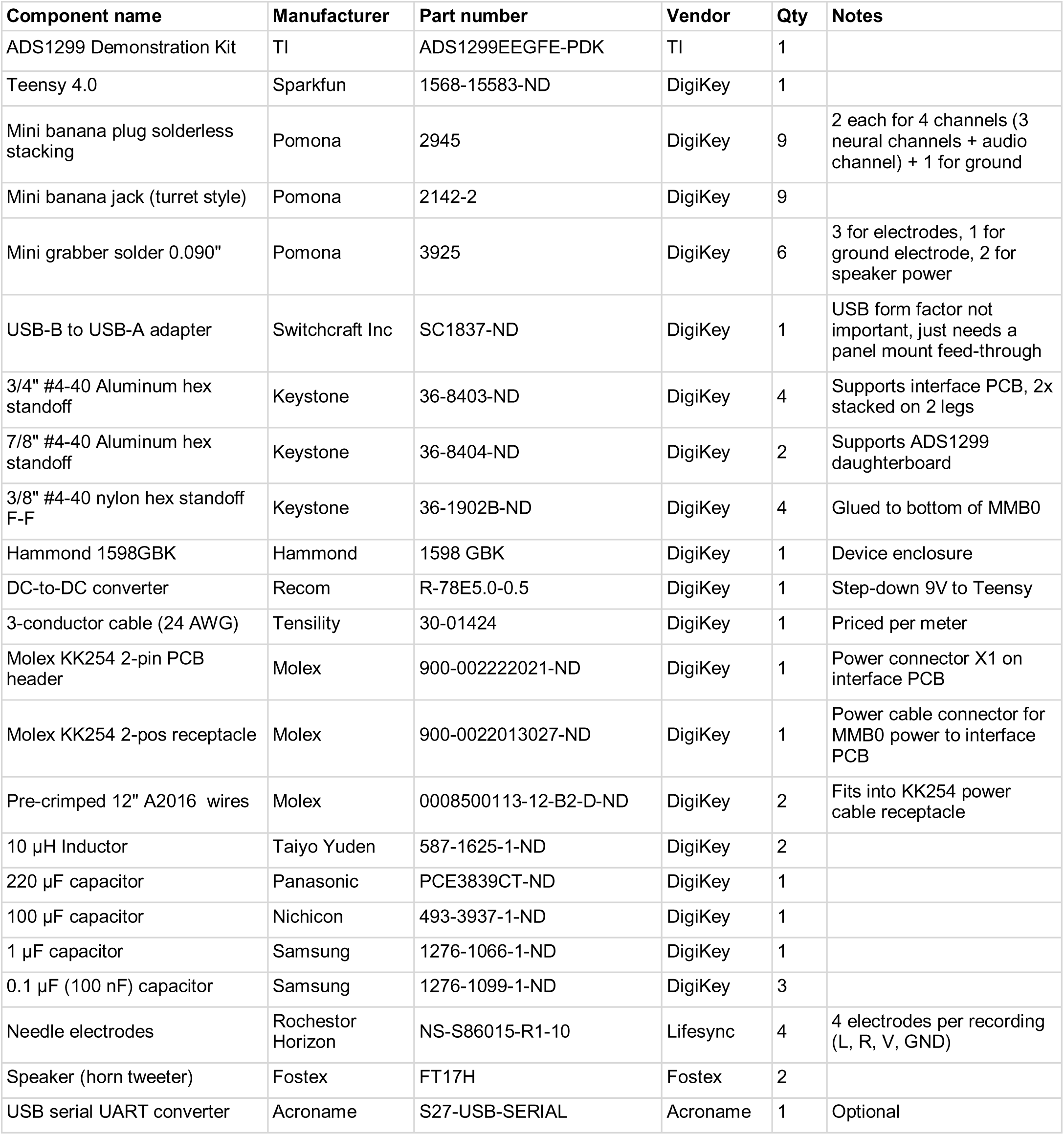
Bill of materials.

A standard desktop PC running Linux communicated with the Teensy over USB. We wrote Python code using the pySerial, multiprocessing, NumPy, Pandas, and SciPy modules to retrieve data over USB and to write data to disk. A graphical user interface (GUI) written in PyQt and PyQtGraph displayed the raw data, filtered data, power spectrum, simultaneous audio input, and computed ABR (***Fig 2***).

**Figure 2.**
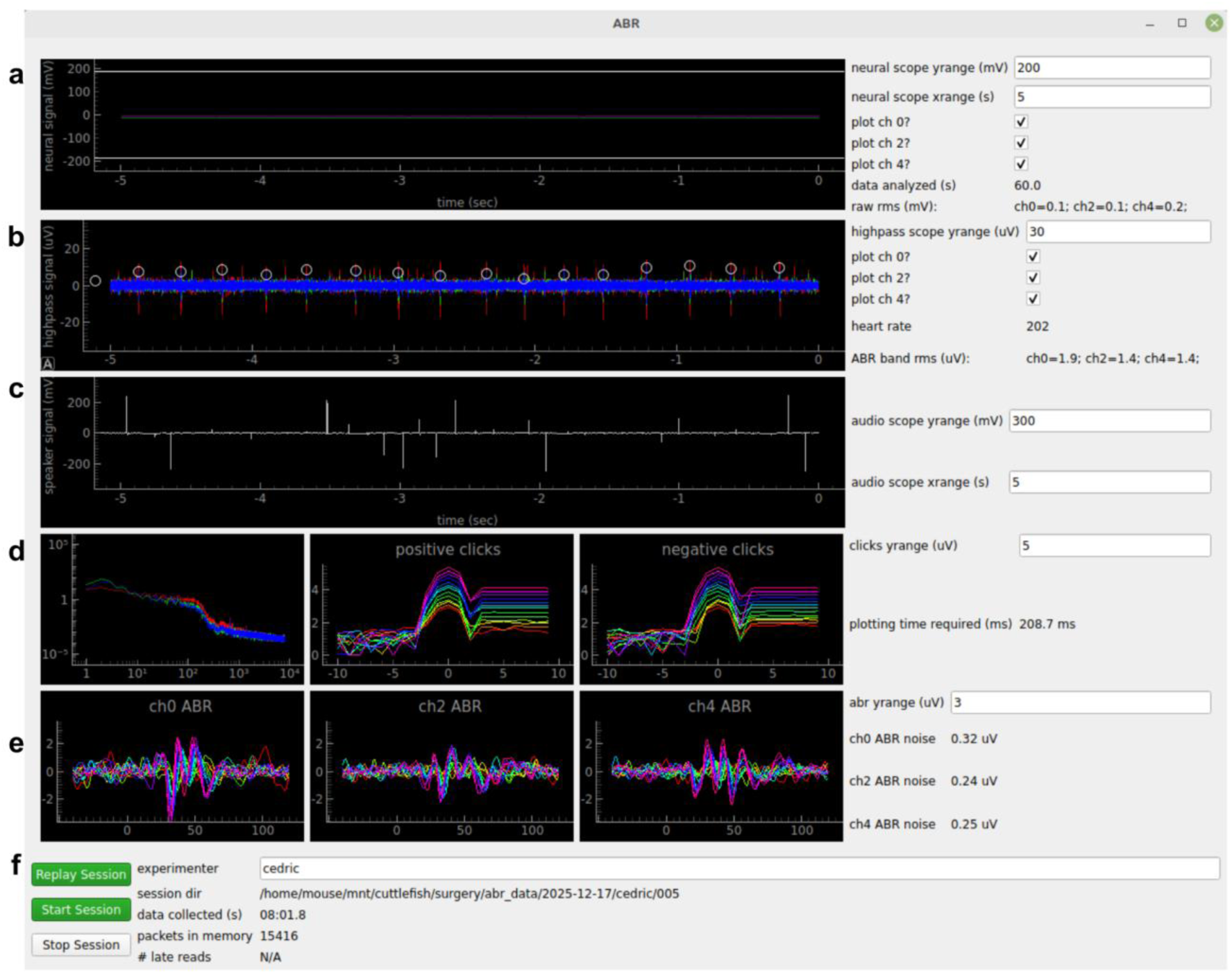
Graphical user interface. ***a)*** Real-time plot of each channel. ***b)*** Same data, filtered in ABR band. *Circles:* detected ECG (heartbeat). ***c)*** Speaker voltage (highpass filtered). ***d)*** *Left:* power spectral density. *Middle and right:* Speaker voltage aligned to sound onset, separated by sound level (colors). ***e)*** Computed ABR. ***f)*** GUI elements to control recordings and view progress.

The total cost of the system was about $400 (excluding the computer), much less than commercially available alternatives.

## Methods

### Subjects

All mice used in this study were first-generation (F1) hybrid offspring of male C57BL/6J and female CBA/CaJ mice. This hybrid cross is resilient to the pronounced age-related hearing loss that afflicts purebred C57BL/6J mice^40,41^. All parent mice were purchased from Jackson Laboratories and all experimental mice were bred in our mouse facility at Emory. Female parent mice were wild-type CBA/CaJ (Jackson Labs 000654). Male parent mice were either wild-type (000664) or 5xFAD mice (034848). The 5xFAD parents carried transgenes related to Alzheimer’s disease and we used them to generate progeny for another study. All progeny from 5xFAD parents were genotyped and no progeny containing 5xFAD mutations were used in this study. Therefore, all mice used in this study were wild-type F1 hybrid C57BL/6J:CBA/CaJ mice.

Mice were raised on a standard (non-reversed) light cycle and were given access to enrichment such as nestlets, a tunnel, chew sticks, and a running wheel. All experiments were conducted on adult mice between 99 and 289 days old. While our study was not designed to detect sex differences, we used males and females in roughly equal proportion. We noted no consistent differences between the sexes except that males were more vulnerable to side effects of dexmedetomidine, as described further below.

### ABR recording procedure

We generally followed standard published protocols^42–44^ for recording the ABR.

*Anesthesia.* Mice were anesthetized during recordings to minimize movement. Anesthesia was induced first by inhalation of 3-5% isoflurane in oxygen using an isoflurane vaporizer (RWD) and an oxygen concentrator (Pureline). Mice were then removed from isoflurane and given an intraperitoneal injection of 100 mg/kg ketamine and 0.8 mg/kg dexmedetomidine or alternatively 100 mg/kg ketamine, 8 mg/kg xylazine, and 1.3-3 mg/kg acepromazine (Patterson Veterinary). We recommend the second anesthetic cocktail because dexmedetomidine can cause severe urinary obstruction in male mice^45,46^ and because we found that including acepromazine led to more reliable anesthetic control. Throughout the recording, mice were administered concentrated oxygen through a silicone facemask (Kent Scientific) at 0.5 L/min. To maintain an adequate level of anesthesia, defined as a negative response to toe pinch, we administered subcutaneous boosters of 30 mg/kg ketamine as necessary, typically about every 30 minutes. In rare cases we delivered additional isoflurane to achieve adequate anesthesia; in those cases we waited to start the recording for at least 1 minute after turning off the isoflurane because of a report that isoflurane can suppress the ABR^13^. After recordings were finished, we injected 2 mg/kg of atipamezole (Patterson Veterinary) to reverse anesthesia and we placed the mouse in a recovery cage on a heating pad until ambulatory.

*Physical setup.* The ABR recording took place in a sound-attenuating chamber that we built from T-slot rail (McMaster) and 1/2” medium density fiberboard (Home Depot). The chamber was cubical with edge length 40 cm. To avoid power line noise from an electrical heating pad, we used a water-recirculating heating pad to warm the anesthetized mouse. This consisted of a heated water pump (Braintree Scientific HTP-1500) connected to a rectangular pad (Össur B-232002402) selected for its small footprint. Body temperature was maintained at 37°C. The mouse’s head was centered between two ultrasonic-capable speakers (Fostex FT17H). The center of the front face of each speaker was 13 cm lateral and 3 cm dorsal to the nearest ear. We defined t=0 as the time at which the speaker began to play the auditory stimulus; no correction factor was included for the time required for sound to reach the ear (about 0.4 ms).

*Electrodes.* We used subdermal needle electrodes (Rochester Horizon NS-S86015-R1-10). We placed the ground electrode on the back near the base of the tail, the ***L*** and ***R*** electrodes ∼2 mm ventral and posterior to the intertragal notch, and the ***V*** electrode within ∼3 mm of the cranial vertex. Each electrode was inserted subcutaneously, parallel to the body, and pointing rostrally.

Good cabling from the electrodes to the ADS1299 was crucial for low-noise operation. We made a custom cable to connect the electrodes to the device’s differential inputs. The electrode end of the cable had three micrograbbers (Pomona 3925-2) connected to shielded three-conductor cable (Tensility 30-01424) approximately 90 cm long, splitting near the ADS1299 into a shielded two-conductor cable for each channel and terminating in banana plugs (Pomona 2945). Using multi-conductor cable ensured that all differential inputs followed the same path for most of their length, and therefore picked up the same electrical noise from their surroundings.

*Audio.* To deliver sound stimuli, we used the software Autopilot^47^ running on a Raspberry Pi 4B with a sound card (HiFiberry Amp2) at a sampling rate of 192 kHz. The sound stimuli were generated as 1-sample clicks, equally likely to be either positive or negative amplitude (acoustic rarefaction or compression) and delivered at an irregular repetition rate of approximately 8 Hz. Each click was delivered at a randomly chosen sound level from 25 to 73 dB SPL at 4 dB intervals.

We used an ultrasonic microphone (Brüel and Kjaer) to measure the produced acoustic waveform, which comprised a sharp peak followed by a brief (∼1 ms) oscillation. Although the acoustic power was concentrated in a 0.5 ms window, we reported the sound level as an average over a time window of 50 ms. (Below 50 ms, perceived volume scales with sound duration^14^, indicating that the integration time of the auditory system is about 50 ms.) Specifically, we divided the maximum peak-to-peak sound pressure by 100 (i.e., 50 ms / 0.5 ms). (We obtained similar results when we explicitly calculated the root mean square amplitude of the recorded sound over a 50 ms window.) Finally, we converted the sound level to decibels (dB) with respect to a 20 μPa reference (dB SPL). The acoustic level of each speaker was approximately equal and approximately linear in amplitude (±2 dB) over the range of sound levels used.

*Experiment design.* Each ABR experiment (i.e., anesthetic session) comprised multiple sequential recordings (n = 18 mice, 37 experiments, 186 recordings). During each recording, the sound was played from either the left or right speaker only. The median recording was 483 s long (range: 37 - 640 s; inter-quartile range: 346 - 501 s). Each experiment contained at least 2 recordings from each speaker side. We excluded recordings with gross artefacts or errors as well as any that differed substantially f rom the other recordings taken in the same experiment, suggesting a shift in electrode placement.

### Induction of hearing loss

In a subset of mice (n=3), we induced hearing loss by bilateral surgical extirpation of the malleus^14,48,49^, one of the ossicles^50–53^ (i.e., middle ear bones). During this surgical procedure, mice were anesthetized with isoflurane delivered through a flexible mask on an infrared heating pad (Kent Scientific). The hair around the ear canal was removed and the skin within the ear canal was disinfected with iodine (Betadine); we avoided chlorhexidine due to the risk of ototoxicity^54,55^. Under a surgical microscope (AmScope), we used fine forceps (Fine Science Tools) to enter the ear canal, break the tympanic membrane, and remove the malleus. Afterwards, the ear canal was flushed with saline. To provide sustained analgesia, 3.25 mg/kg extended-release buprenorphine (Patterson Veterinary Ethiqa XR) was administered peri-operatively. As a control, other mice received sham surgeries that were identical except that the tympanic membrane and malleus were left intact.

Mice were randomly assigned to either bilateral or sham hearing loss surgery. The experimenter collecting and analyzing ABR data was blinded to their condition. We recorded ABR in two pre-surgery sessions and two post-surgery sessions. The two pre-surgery sessions occurred 3-7 days and 0-1 days before the surgery. The two post-surgery sessions occurred 3 days and 5-6 days after surgery.

### Data analysis

All data analysis was conducted in Python using the packages IPython^56^, Pandas^57^, NumPy^58^, SciPy^59^, statsmodels^60^, and matplotlib^61^. We reported statistical significance as p > 0.05 (not significant), p < 0.05 (*), p < 0.01 (**), or p < 0.001 (***).

## Results

### High-fidelity measurements of the ABR

We implemented a standard signal processing pipeline to compute the ABR (***Fig 3a***), beginning with spectral filtering. The power spectrum of the raw data declined with frequency, in approximate accord with the 1/f power law that describes many biological signals^38^ (***Fig 3b***). Raw ABR recordings contained large (∼100 μV) artefacts repeating about once per second, which we attribute to electromyographic (EMG) signals or movement artefacts induced by breathing (***Fig 3c, upper***). To isolate the desired signal from these artefacts, we used a 2nd order Butterworth bandpass filter of 300 - 3000 Hz, which we called the “ABR band”. The amplitude of the signal within the ABR band was about 1 - 1.5 μV rms (root mean square). Within this band we observed large-amplitude (∼30 μV) spikes repeating around 3-4 Hz (***Fig 3c, lower***). Based on the waveform shape (***Fig 3d,e***) and periodicity, we attribute these spikes to the electrocardiogram (ECG) from the heart^8,62^. We took advantage of the reliable ECG signal to validate that the electrodes were in contact and to display heart rate as an indicator of anesthetic state in our software.

**Figure 3.**
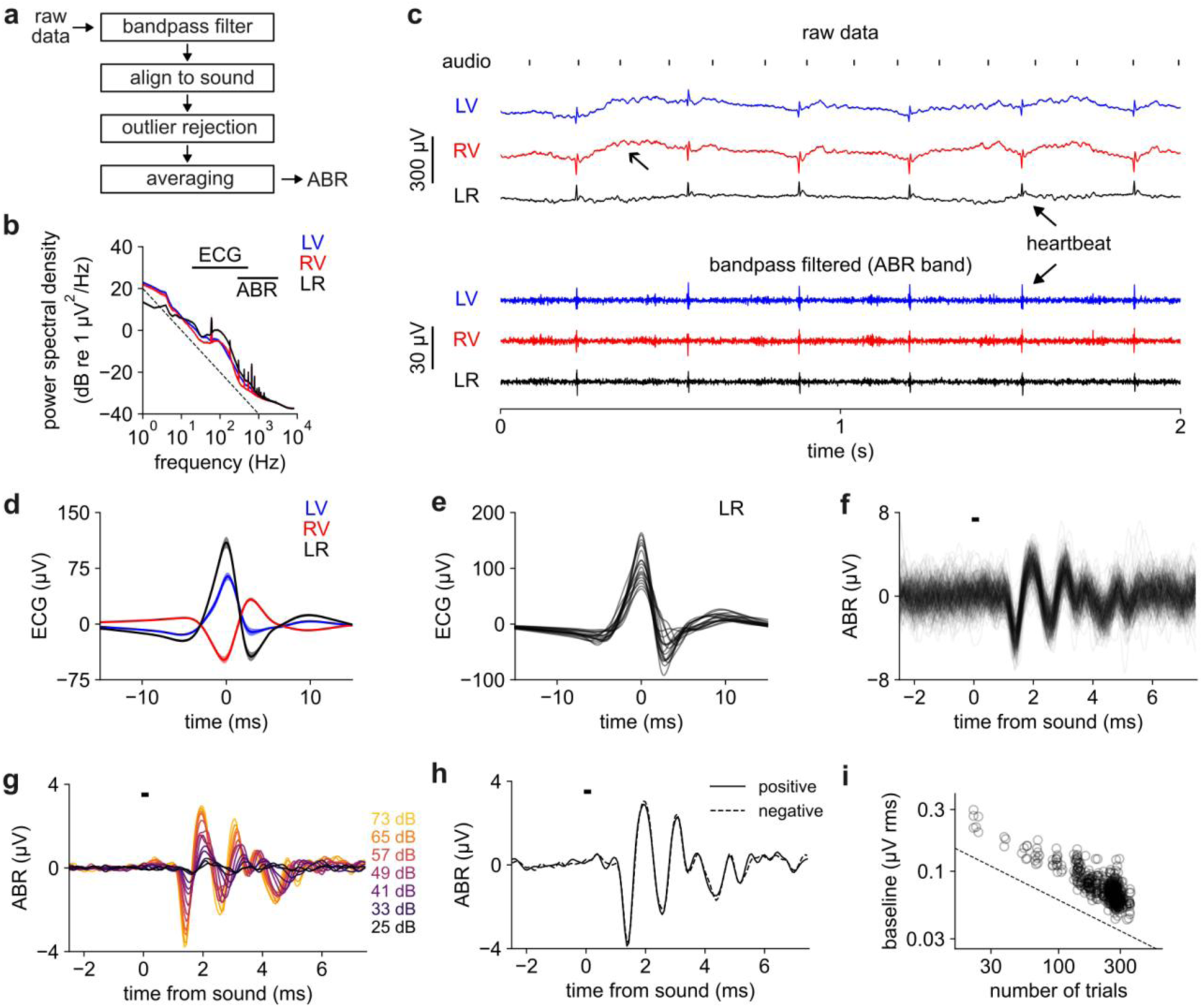
Signal processing for ABR estimation. ***a)*** Signal processing pipeline. ***b)*** Power spectral density of the raw data on each channel. Error bars: SEM over n = 18 mice. Horizontal lines: ECG and ABR bands. Diagonal dashed line: 1/f slope. ***c)*** *Top:* 2 s of raw data (baseline removed with 0.1 Hz highpass filter). *Bottom*: data filtered in ABR band (300-3000 Hz). Open arrow: putative breathing artefact. Closed arrows: ECG (heartbeat). ***d)*** ECG filtered (20-500 Hz) and aligned to peak. Error bars: SEM over n = 18 mice. ***e)*** ECG on LR channel. Separate line for each mouse. ***f)*** Multiple individual trials of ABR evoked by the loudest sound level. Horizontal bar: click stimulus. All data in panels ***f***-***h*** comes from one example recording (***RV*** channel in response to left click). ***g)*** ABR grouped by sound level (color). ***h)*** ABR grouped by positive or negative click. ***i)*** Baseline (t = -1.875 ms) of trial-averaged ABR declines with square root of the number of trials. Circles: 186 recordings with 3 channels each (558 points total) from 18 mice. Dashed line: slope of n^-1/2^.

After spectral filtering we temporally divided the data into individual trials defined as 2.5 ms before and 7.5 ms after each sound stimulus (i.e., click from left or right). Because the ECG overlapped the ABR band (***Fig 3b***) it could not be removed through spectral filtering, but its short duration and large amplitude meant that it could readily be removed through outlier rejection. Specifically, we dropped trials whose absolute value or rms level exceeded the typical trial by at least three standard deviations. After outlier rejection, the ABR to the highest sound level was partially visible on individual trials without averaging (***Fig 3f***), which surprised us given the small size of the signal and which demonstrated the high signal-to-noise ratio of this system. Finally, we computed the ABR by averaging together all the trials of the same sound level within a recording (***Fig 3g***), revealing lower-amplitude ABRs below the single-trial noise floor.

To assess electrical stimulus artefact from the speaker, we measured the ABR to positive and negative clicks (rarefaction vs compression), which would be expected to elicit opposite electrical artefacts but similar physiological response (***Fig 3h***). The response to both polarities was quite similar, indicating that minimal stimulus artefact was included.

Increasing the number of included trials should decrease the noise level in the averaged ABR by the law of large numbers. Indeed, we found that the rms level during the baseline period (-2.5 to -1.25 ms from click onset) declined in inverse proportion to the square root of the number of included trials (***Fig 3i***), as expected for a Gaussian random variable. Our recordings typically contained 100 - 300 trials per sound level, resulting in about 0.05 - 0.15 μV rms noise in the trial-averaged ABR baseline. This noise level set the minimum detectable ABR size; increasing the trial count would decrease this noise floor further.

### Consistency and variability in the ABR

On all three channels (***LV***, ***RV***, ***LR***) and in response to clicks from either side, the ABR exhibited a characteristic oscillation with about a 1 ms period (***Fig 4a***; grand average over n = 18 mice). Compared to higher sound levels, lower sound levels evoked a response of similar shape but smaller and slightly delayed, which could be readily visualized in a heatmap representation (***Fig 4b***). By cross-correlating the response across different sound levels (***Fig 4c***) we estimated that the delay increased at lower levels with a slope of 6.9 ± 0.8 μs/dB (standard deviation over mice). Burkard *et al.* reported a steeper slope in rats and proposed that steeper slopes were correlated with larger head size across species^14^.

**Figure 4.**
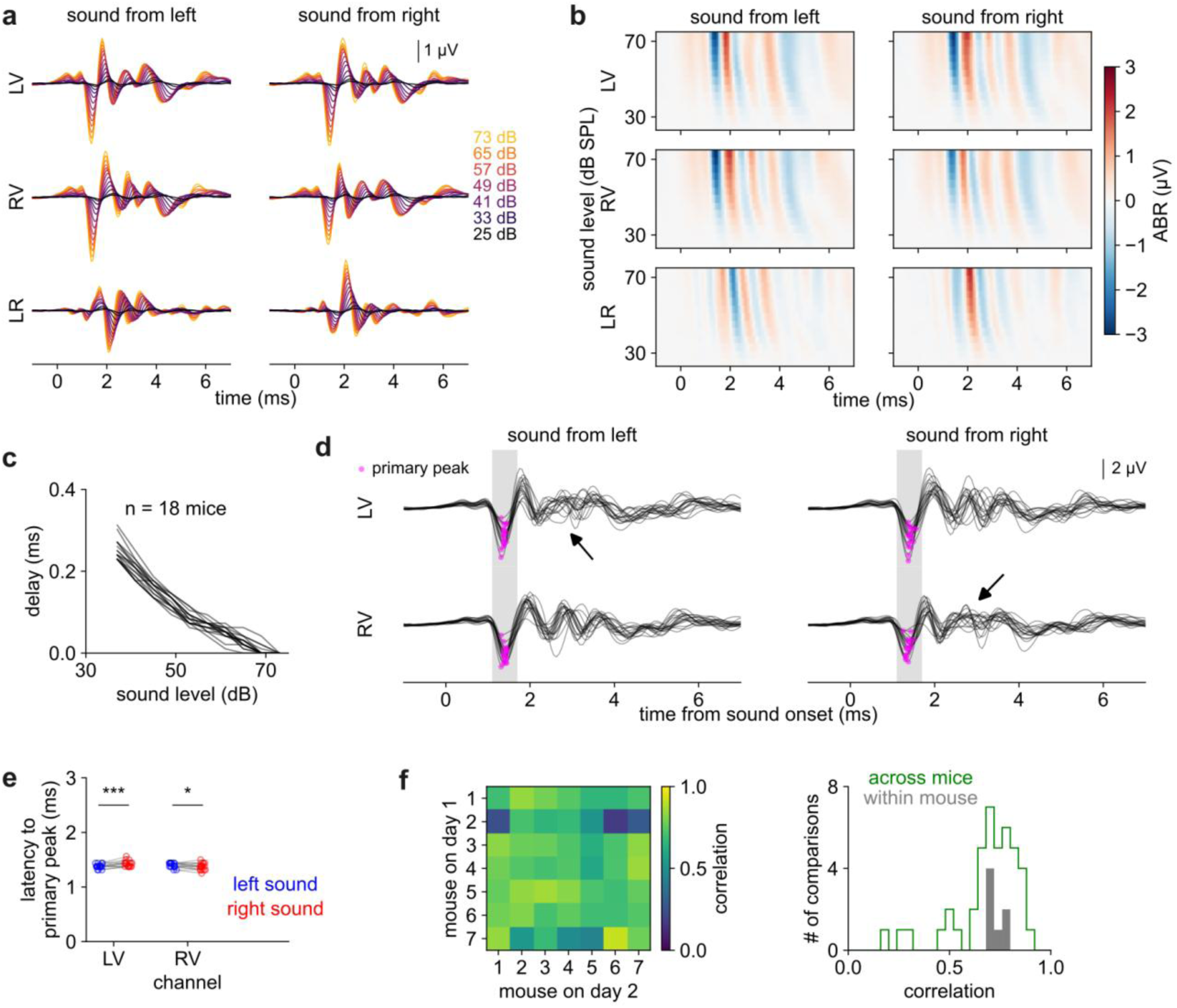
Consistency and variability in the ABR. ***a)*** ABR by sound level (colors), channel (rows), and speaker side (columns) averaged over n = 18 mice. ***b)*** Heatmap of data in panel (a). ***c)*** The ABR is increasingly delayed at lower sound levels by 6.9 μs / dB (linear fit) relative to the response at the highest tested level. Individual lines: n = 18 mice, each averaged over session, channel, and speaker side. ***d)*** Overlaid ABR from n = 18 mice (loudest level only). Magenta: primary peak. Later peaks are less consistent, especially for ipsilateral sounds (black arrows). ***e)*** Latency of primary peak. Ipsilateral sounds evoked significantly shorter latencies (paired t-test). ***f)*** *Left:* Correlation (Pearson’s r) between ABR taken on day 1 (rows) and day 2 (columns) over all pairs of n = 7 mice. Values along the diagonal are within-mouse; off-diagonal values are across-mouse. *Right*: Distribution of within-mouse (gray) and across-mouse (green) correlations.

While the ABR was somewhat variable across mice, the most reliable feature was a large, early, negative peak, which we called the “primary peak” (magenta dots in ***Fig 4d***). The primary peak occurred at 1.393 ± 0.010 ms after sound onset (SEM over n = 18 mice) and was slightly earlier for ipsilateral (1.375 ms) than for contralateral (1.411 ms) clicks, consistent with the expected ∼0.03 ms interval for sound to travel the ∼1 cm between the ears. Keeping in mind that we always placed the positive (non-inverting) electrode on the ear (as described in Methods), the negativity of the peak indicated neuronal activation near the ear at this latency^38^. Based on the large size and consistent timing of this peak (***Fig 4e***), we attribute it to what prior work^1^ called “Wave 1”, which likely originates from the afferent auditory nerve. Later peaks were more variable in their timing and could even be absent, especially for ipsilateral sound (black arrows in ***Fig 4d***), and so we did not pick them out or number them.

We asked whether each mouse had an idiosyncratic “signature” ABR, which might differ across mice but be consistent over sessions from the same mouse. Alternatively, the ABR might vary across sessions even within the same mouse. To distinguish these possibilities, we recorded the ABR in a second session from a subset of mice (n = 7) and computed the correlation both between mice and also across sessions from the same mouse. We found that correlation across sessions from the same mouse (0.73 ± 0.03) was similar to the correlation between mice (0.69 ± 0.16, ***Fig 4f***). Some pairs of mice showed especially low or high correlation relative to the fairly narrow range of within-mouse correlations. Overall, we found little or no evidence that each mouse had an idiosyncratic signature ABR.

### Differences in LV and RV channels produced sensitivity to sound direction in the LR channel

The auditory system localizes sound sources by computing differences between the left and right ear^16,63,64^. While many previous studies have recorded the ABR on one side only (generally ipsilateral to the sound), our experiments measured the ABR on both sides of the head, allowing us to measure inter-hemispheric differences directly.

In the vertex-ear channels ***LV*** and ***RV***, contralateral and ipsilateral clicks at the loudest level evoked a primary peak of similar magnitude (***Fig 5a***). The similar response to contralateral and ipsilateral sound was surprising because we expected contralateral responses to be smaller, due to attenuation of sound as it passed through the head (i.e., acoustic shadowing). Moreover, the primary peak was so early (<2 ms) that contributions from the contralateral hemisphere should be minimal^8^. Instead, the primary peak in response to contralateral sound appeared to be, if anything, slightly larger than the primary peak in response to ipsilateral sound.

**Figure 5.**
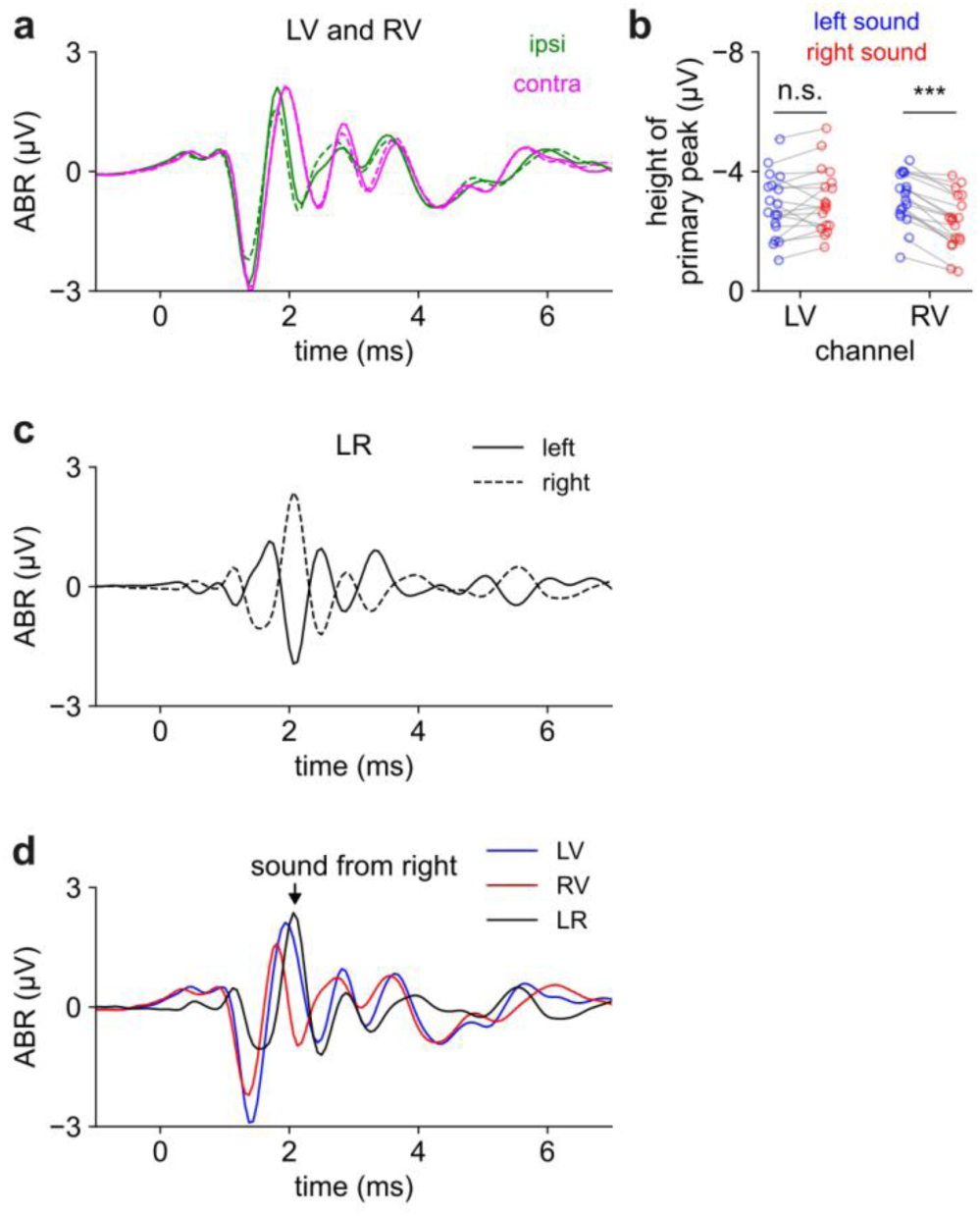
The ABR is sensitive to sound direction. ***a)*** Ipsilateral (green) and contralateral (magenta) sounds evoke a similar early response (t<2 ms) before diverging later. Results are similar for sound from left (solid) or right (dashed). Panels A, C, and D show a grand average from n = 18 mice. ***b)*** The magnitude of the primary peak is similar for ipsilateral and contralateral sounds with a slight bias toward contralateral sound. ANOVA described in text. Asterisks here indicate post-hoc t-tests (n = 18 mice). ***c)*** The ***LR*** channel differentiates left (solid) vs right (dashed) sound, especially around 2.1 ms. ***d)*** All channels overlaid for sound from the right. ***LR*** channel is maximal when ***LV*** and ***RV*** channels diverge (arrowhead).

We statistically quantified the amplitude of the primary peak using a 2-way ANOVA with main effects of channel (***LV*** or ***RV***), speaker side (L or R), and an interaction term (contralateral or ipsilateral). All three terms were statistically significant to varying extents: a larger primary peak was associated with contralateral sound (interaction term: p < .001; effect size β = 0.47 µV), sound from the left (p < 0.05; β = 0.29 µV), and recording on the LV channel (p < 0.05; β = 0.24 µV). Due to the combination of these factors, contralateral responses were stronger on the RV channel (p < 0.001; post-hoc t-test) but not on the LV channel (p > 0.05; ***Fig 5b***). Notwithstanding these relatively small differences, the primary peak was notably similar in magnitude across all conditions; none of the effects noted above amounted to more than 15% of the size of the typical primary peak (3.45 µV).

The contralateral and ipsilateral responses diverged more strongly after the primary peak, an effect that was most apparent in the ***LR*** channel since by definition it was the difference between ***LV*** and ***RV***. The ***LR*** channel peaked at 2.097 ± 0.018 ms after left clicks and 2.083 ± 0.014 after right clicks (SEM over n = 18 mice; ***Fig 5c, 5d***). Unlike ***LV*** and ***RV***, the ***LR*** channel was highly sensitive to sound direction: this peak was negative in response to sound from the left but positive in response to sound from the right, likely reflecting stronger neuronal activation on the side ipsilateral to the speaker at this timepoint.

### A peak-agnostic response measure: rolling standard deviation

After the primary peak, other peaks in the ABR waveform were somewhat inconsistent across mice, especially for ipsilateral sound (***Fig 4d***). Moreover, we found peak-picking algorithms to lack robustness because they were sensitive to noise and required fine tuning of multiple inter-dependent parameters to pick up subtle peaks at low levels while rejecting false peaks and shoulders. Indeed, previous work has reported surprising variability in the peaks picked by human observers, even highly trained ones^9^. Thus, we sought a peak-agnostic measurement of response strength.

We defined the “response strength” of the ABR as its rolling standard deviation over time. That is, at any particular timepoint the response strength was the standard deviation of the ABR over a 1.25 ms window of data centered on that timepoint (***Fig 6a***). When we needed to define a single (scalar) response strength, we used the response strength at t = 2.0 ms after sound onset, which was when the rolling standard deviation tended to peak. While the response strength increased roughly linearly with sound level, its variability scaled with its strength (i.e., its coefficient of variation was constant). Therefore, we generally plotted response strength on a logarithmic axis, resulting in a more consistent variance across sound level (***Fig 6b***).

**Figure 6.**
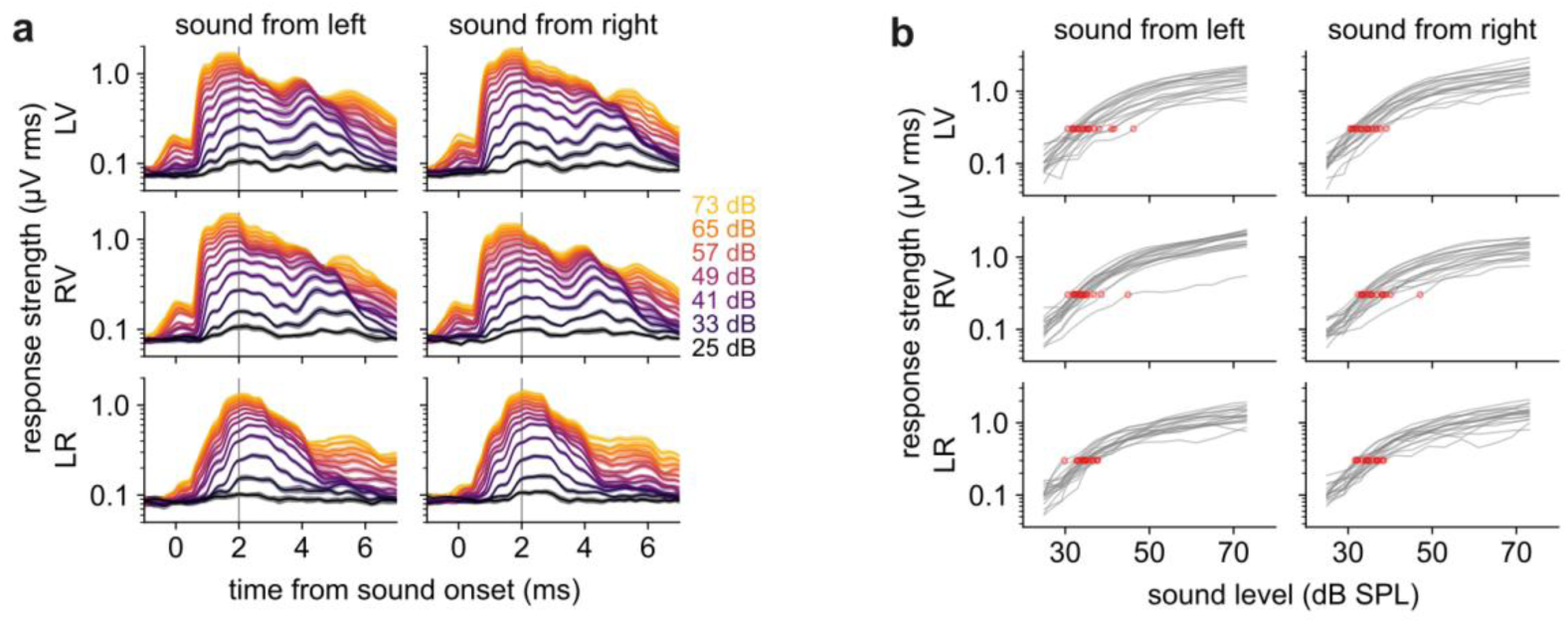
ABR strength increases with sound level. ***a)*** Response strength separated by sound level (colors) using a rolling window of 1.25 ms. Shaded error bars: SEM over n = 18 mice. Peak typically at 2 ms (gray vertical line). ***b)*** Response strength (t=2.0 ms post-click) versus sound level, separated by channel (rows) and speaker side (columns). Gray lines: individual mice. Red dot: threshold.

Auditory sensitivity is typically measured as the threshold level, or minimum level of sound necessary to evoke a detectable ABR^1^. However, detectability could depend as much on experimental parameters like noise level and trial count as on the strength of the ABR signal. In our work, we defined the threshold level as that level which elicited an ABR strength exceeding 0.3 μV (***Fig 6b***). We chose 0.3 μV because it was above the baseline response (defined as the ABR strength 1.875 ms before sound onset) over 99% of the time, meaning false positives were rare. Moreover, ABR strength steeply increased with sound level around this value (***Fig 6b***), enhancing our ability to detect changes in threshold level. To estimate threshold precisely, we linearly interpolated the response strength between the sound levels we tested.

Using these metrics, the mean threshold level across healthy-hearing mice was 34.9 ± 2.3 dB SPL (standard deviation over n = 18 mice). Thresholds were similar across ***LV***, ***RV***, and ***LR*** channels (34.7, 35.3, 34.8). In ***LV*** and ***RV***, thresholds were slightly lower for contralateral (34.1) than for ipsilateral (35.9) sound, consistent with the finding in the previous section that the primary peak was slightly larger for contralateral sound. We note that differences in threshold within a study are more interpretable than its exact value, given variation in experimental conditions and analytical parameters across studies.

### The system is capable of detecting threshold changes after bilateral hearing loss

Finally, we evaluated the ability of this system to detect changes after hearing loss, a common usage of ABR. We induced bilateral hearing loss in n = 3 mice by surgical extirpation of the malleus, a middle ear bone that enhances the transmission of acoustic energy to the inner ear. Because we left the inner ear intact, this manipulation should produce only a partial deafening^14,48,49^. As a control, we performed sham surgery on an additional n = 3 mice, leaving both ears intact. In all mice, we measured the ABR before and after the surgery. The first post-hearing loss measurement was taken 3 days after surgery. See Methods for additional details.

Qualitatively, the ABR was unaffected by sham surgery but greatly attenuated by bilateral hearing loss (***Fig 7a,b***), across all three channels and in response to sound from both sides. In mice with bilateral hearing loss, we observed no detectable ABR at the lowest sound levels and an attenuated response at higher sound levels (***Fig 7c***), suggestive of a rightward shift (i.e., increased threshold) rather than a downward shift (i.e., overall decreased response). The threshold increased by 25.5 ± 3.3 dB after bilateral hearing loss (standard deviation over n = 3 mice) while sham surgery had no effect (-0.3 ± 1.5 dB; ***Fig 7d***). Notably, the responses at the highest sound levels were nearly the same in bilateral and sham mice, indicating that the auditory system retained sensitivity to loud sound and suggesting that a compensatory process steepened the response curve. Taken together, these data show that our system is capable of measuring a consistent threshold shift induced by hearing loss.

**Figure 7.**
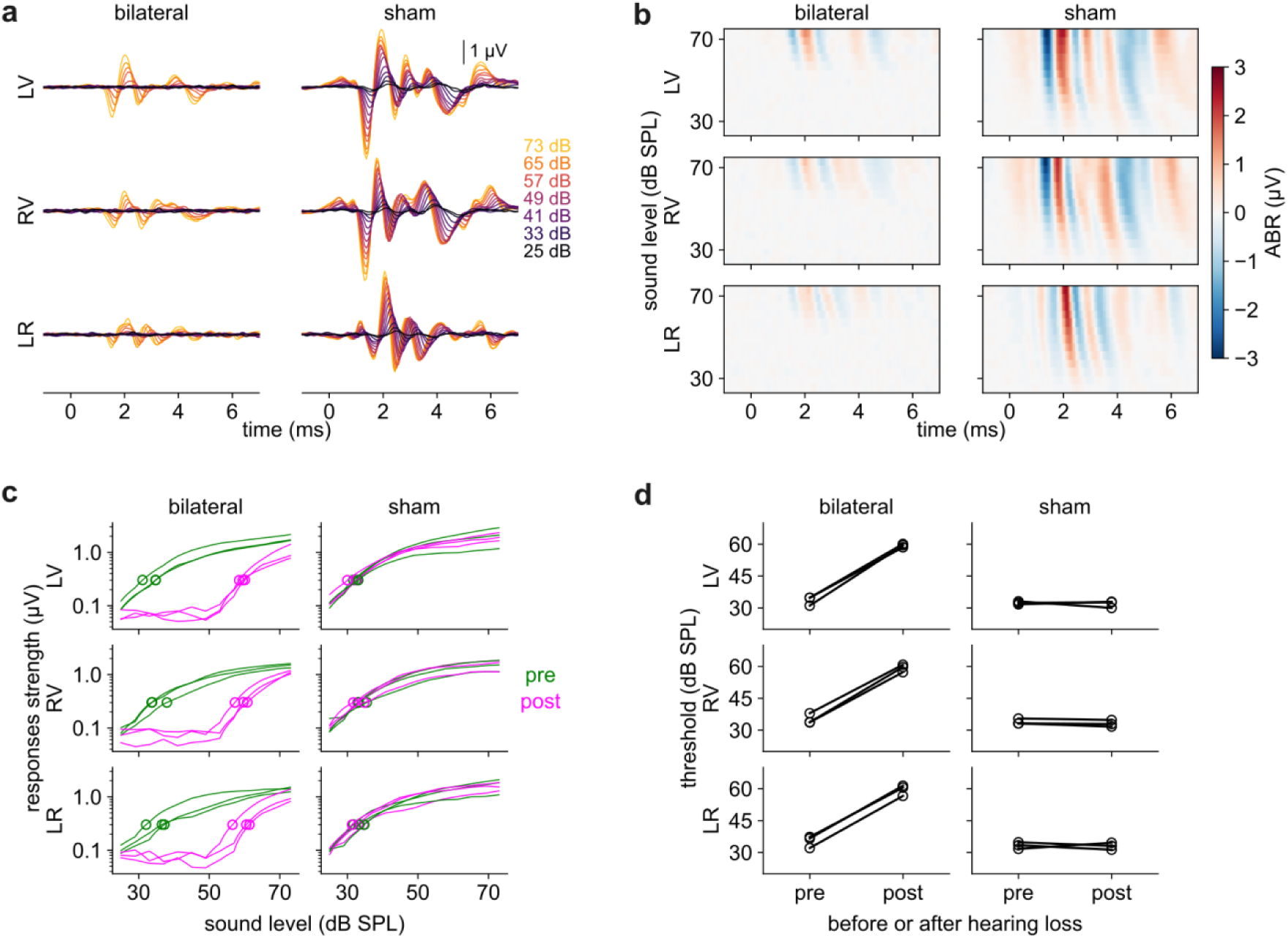
Detecting hearing loss. ***a)*** ABR is attenuated after bilateral (left column; n = 3 mice) but not sham (right column; n = 3 mice) hearing loss. This figure shows data for right sounds only; left sounds elicited similar results. ***b)*** Same data as panel (***a***). ***c)*** Response strength versus level (green: pre-surgery; magenta: post-surgery) is shifted rightward after bilateral hearing loss (left column) but not sham surgery (right column). ***d)*** Threshold elevated by 25.5 ± 3.3 dB (standard deviation) after bilateral hearing loss (left column). No effect (-0.3 ± 0.9 dB) after sham hearing loss (right column).

## Discussion

We designed an open-source system for multi-channel recording of the ABR with high fidelity. We measured the ABR in three differential channels (***LV***, ***RV***, and ***LR***) and quantified the timing, magnitude, and variability of the response. As expected, we found that the ABR was attenuated and delayed with decreasing sound level. The ***LR*** channel emphasized differences between the two sides and was sensitive to sound direction. The ***LV*** and ***RV*** channels exhibited a consistent “primary peak” around 1.4 ms likely originating from the auditory nerve; surprisingly, this peak was no greater for ipsilateral sound and in some cases it was greater for contralateral sound. Multiple recordings from the same mouse were no more similar than were recordings from different mice, indicating little evidence for a “signature” ABR for each individual. We calculated the threshold of auditory sensitivity using a peak-agnostic measurement of response strength (rolling standard deviation) and we showed that we could detect an increased threshold after surgical extirpation of the malleus. In summary, this system is capable of sensitive measurements and is suitable for detecting hearing loss in mice. Moreover, we measured the ABR, electrocardiogram (ECG), and electromyography (EMG) with the same system, indicating that it could likely be used for non- or minimally-invasive measurements of other biosignals like the auditory steady-state response (ASSR), visual or somatosensory evoked potentials (VEPs or SEPs), or 8-channel electroencephalography (EEG).

### Challenges encountered and solutions employed

Because the ABR is such a small signal, it is important to minimize noise. Our pilot recordings were contaminated by electrical artefact from low-quality speakers (data not shown). Alternating positive and negative stimuli was a simple and effective way to cancel out this artefact; ultimately, we switched to speakers that produced less electrical artefact, perhaps because they better matched the impedance of our audio amplifier. Another source of noise in pilot recordings was mismatch between the length and path of the cables from each electrode, which caused them to pick up noise differently. We addressed this by fabricating a single multi-conductor shielded cable. Finally, the use of an electric heating pad introduced substantial noise in pilot recordings, which we addressed by switching to a water-based heating pad. Together, these changes eliminated all substantial sources of noise and enabled high-quality recordings.

It is important to ensure that the electrodes are placed consistently across recordings and that they do not slip out during the experiment. It is plausible that variability between the waveforms we recorded arose in part from subtle changes in electrode placement^12^. Adhesive skin electrodes may provide more consistent (but possibly attenuated) recordings and would also be less invasive.

### Limitations of our system and plans for future development

We used a relatively simple stimulus: a click from left or right at various amplitudes. The system could be extended to more complex stimulus sets by using one of the digital inputs on the Teensy to detect a digital code indicating the timing and identity of sound played on that trial.

The ADS1299 contains additional useful functionality, such as impedance checking and “lead-off” detection (i.e., when the electrodes lose contact with the subject). Moreover, this chip is capable of using an active reference electrode to cancel out noise with a real-time feedback loop. These functions could be incorporated into a future version of the system. We invite potential users to consider customizing the system for their own applications in biosignal measurement.

### The effect of bilateral conductive hearing loss

We found that bilateral hearing loss induced a moderate but consistent threshold shift of 25.5 ± 3.3 dB (standard deviation). This threshold shift is relatively small—less than the attenuation of over-the-counter foam earplugs—and less than the 30-50 dB threshold shift reported in gerbils^48,49^ and 50 dB reported in rats^14^ after the same surgical procedure, which may indicate inter-species differences in the role of the middle ear bones in amplifying sound^50,51^.

We recently reported that bilateral hearing loss induced with the same surgical procedure robustly impaired the ability of mice to perform a sound-seeking task at sound levels well above threshold. Surprisingly, unilateral hearing loss resulted in strong but transient impairment in sound-seeking, before a notable and near-complete recovery^65^. To better explain the behavioral effects and subjective experience of hearing loss, future studies will need to make additional physiological measurements beyond threshold shifts.

### Vertex-ear vs between-ear ABRs

Relatively few studies have compared the ABR on both sides of the head to both left and right sounds. At the earlier timepoints before 2 ms, we found that responses to ipsilateral and contralateral free-field sound were fairly similar. While we expected that the contralateral response would be delayed, it was surprising that the early contralateral response was slightly larger than the ipsilateral response, given that the early response is generally attributed to the auditory afferent nerve (which does not cross the midline). This unexpected result may be due to the presence of echoes, other connections that do cross the midline, or adaptation to our experimental design of blocks of stimuli presented from the same side. Future experiments with in-ear delivery of interdigitated left and right sounds may differentiate between these possibilities.

In all channels, ipsilateral and contralateral responses diverged at later timepoints (2-3 ms). The ***LR*** channel (that is, the pinna-to-pinna electrode montage) was sensitive to the differences between the left and right side of the head and may be particularly useful for studying spatial tuning and unilateral hearing loss^6^.

### Challenges in interpreting the ABR

In a comprehensive handbook for practitioners, Hall discussed what we call the “anatomical single origin” hypothesis: that the different waves of the ABR correspond in a one-to-one fashion to different brain structures (e.g., Wave 2 arises from the cochlear nucleus, Wave 3 from the olive, and so on). This hypothesis is predicated on the assumption of time-locked serial activation of those structures (a “synfire chain”), which appears somewhat unlikely given the prominent role of crossover and feedback projections throughout multiple levels of the auditory system. The single origin hypothesis would also require that individual brain structures cause individual extracranial voltage peaks, but the mapping from intracranial activity to extracranial voltage is in fact quite complex^38^ and will depend on the anatomical shapes of these structures, which differ between species^8,12^. A rigorous test of this hypothesis would require biophysical modeling and recordings before and after ablation of each structure, a comprehensive approach which has rarely been attempted except for one classic series of studies in cats from Melcher *et al.*^20–22^.

Hall ultimately dismissed the anatomical single origin hypothesis as “*clinically appealing yet overly simplistic … inaccurate … [and] erroneous*”. Hall made an exception for “*the unequivocably accurate statement that ABR wave I arises from the auditory nerve*”, but one possible explanation for our results would be that midline-crossing connections might in fact contribute to the primary peak (although see refs^8,15^). Moreover, our study found no more similarity between recordings from the same mouse on different days than between different mice, suggesting that the shape of the ABR waveform either varies over time or strongly depends on measurement conditions like electrode placement^12^.

For all of the above reasons, we would suggest caution in ascribing neurological meaning to individual ABR peaks. We suggest instead that the ABR may be a brief wavelet-like oscillation generated by the auditory system as a whole, with relatively greater contribution from the periphery to the earlier part of the oscillation^66^. Specifically, we suggest a peak-agnostic measurement of ABR strength based on the rolling standard deviation.

Although the ABR can indicate peripheral hearing loss, it is also driven by central auditory brain regions, some of which send efferent output to the cochlea. Reflecting this complexity, some people with abnormal ABR due to auditory neuropathy demonstrate normal hearing on some tests^4^ while some people with normal ABR report disturbed hearing (“hidden hearing loss”)^6^. Central auditory plasticity may be either adaptive or maladaptive^4,67^. Thus, while the ABR is a useful tool for measuring some physiological properties of the auditory system, it is not a “hearing test” and must be complemented with behavioral or cognitive measures in order to better capture the auditory experience.

## Author Contributions

RG developed the experimental approach and performed most of the recordings. CB performed additional recordings. AFM performed all hearing loss surgeries for the dataset included in the manuscript and JM performed hearing loss surgeries for pilot experiments. WNG developed the auxiliary hardware and the Teensy software to interact with the ADS1299, designed the physical enclosure, and developed an early version of the desktop software. RG and CCR analyzed the results and wrote the paper. CCR conceived of the project and developed the final version of the desktop software. All authors approved the final version of the manuscript.

## Acknowledgements

We would like to acknowledge Jonathan Newman for suggesting the overall approach and the ADS1299, Dan Polley for guidance on the choice of hearing loss procedure and best practices for ABR, Julia Lazo and Danielle Babbitt for guidance on the effects of dexmedetomidine in male mice, and Vaibhav Thakur and Nick Steinmetz for guidance on sound-attenuating chamber construction.

Research reported in this document was supported by the National Institute Of Deafness and Communication Disorders under Award Number R21DC019711 and Award Number F31DC022802 (supporting McElroy); the National Institute Of Neurological Disorders And Stroke of the National Institutes of Health under Award Number R34NS137017; the National Institute Of Aging of the National Institutes of Health under Award Number T32AG087922 (supporting Bowe) and under Award Number U24AG072122 via the Alzheimer’s Association’s and the National Alzheimer’s Coordination Center’s New Investigator Award Program. The content is solely the responsibility of the authors and does not necessarily represent the official views of the National Institutes of Health.

For the 5xFAD breeder mice, we acknowledge Robert Vassar, PhD and the Mutant Mouse Resource and Research Center (MMRRC), which is supported by the National Institutes of Health.

Additional financial support was provided by a Whitehall Foundation Research Grant, the Center for Neurodegenerative Disease at Emory University, and an Emory University undergraduate research fellowship (supporting JM).

## Assurances

The authors declare no competing interests.

All animal experiments were conducted under the guidance and approval of the Emory University Institutional Animal Care and Use Committee (IACUC).

All data presented here may be downloaded from Zenodo^36^ at https://doi.org/10.5281/zenodo.17933111. The Teensy and Python code used to run the experiments, the hardware schematics used to build the system, and the Python code used to generate the figures are all freely available on our github repository at https://github.com/Rodgers-PAC-Lab/ABR2025.

