## Supplemental Material for "An open-source system for recording the auditory brainstem response (ABR) and an example application in mice with hearing loss"

### Supplemental Information for Gargiullo 2025

#### Supplemental Methods

This section provides further information on how to construct and use the system that we designed for measuring auditory brainstem response (ABR).

##### Overview of the system

Excluding electrodes, cabling, and the desktop PC, the entire system is contained within a plastic enclosure box (***Supplemental Fig 1***). The electrodes were connected to the enclosure's front panel and the desktop PC connected to a USB port on the enclosure's back panel. The enclosure contained an MMB0 motherboard, an ADS1299 daughterboard, and a custom-designed interface PCB. In turn, the interface PCB housed a Teensy 4.0 microcontroller, a voltage regulator, and a USB debugger. The MMB0 provided power to the daughterboard and to the voltage regulator on the interface PCB, which in turn powered the Teensy and the USB debugger.

The MMB0, daughterboard, Teensy, enclosure, voltage regulator components, and miscellaneous components are all commercially available (see Bill of Materials). We provide the design files for the interface PCB on our github (https://github.com/Rodgers-PAC-Lab/ABR2025), which can be printed by a vendor such as OSH Park.

The Teensy was connected to all other components (***Supplemental Fig 2***):

- It exchanged data with the ADS1299 using a 4-wire SPI protocol from a header on the interface PCB labeled 'extra' to a header on the daughterboard labeled J3.
- It exchanged data with the desktop PC over its USB connection.
- It controlled LEDs labeled "power" and "data" on the front panel
- It could be connected to an optional USB debugger chip, which could be connected to the desktop PC through a second USB cable.

##### Construction of the system

*Physical enclosure.* We drilled holes in the enclosure's front panel to accommodate the power switch, banana jacks, and LED indicator lights. In the rear panel we drilled a hole to provide access to the MMB0's DC-IN jack and one to accommodate the USB-B to USB-A adapter.

*Teensy 4.0.* Normally, the Teensy would be powered by the USB connection, but we wanted it to be powered instead by the MMB0 motherboard so that it could be powered off and on by the front panel. To prevent contention on these power lines, it is necessary to cut the trace between the VIN and VUSB pads on the back of the Teensy (***Supplemental Fig 3a***).

To provide connections to other devices we soldered male pin headers for all through-hole pins, as well as the surface mount pins 24-33 on the bottom of the Teensy.

*MMB0 motherboard.* First, we soldered a power cable to the positive and ground leads of the MMB0's DC-IN jack (***Supplemental Fig 3b***). The other end of this cable connected to the interface PCB.

Second, to insert a power switch into the circuit, we clipped the rear positive leg of the DC-in jack where it connects to the PCB (***Supplemental Fig 3c***). We wired this clipped leg of the DC-IN jack to the positive leg of the power switch, then connected the negative leg of the switch to the jack's ground pin.

Third, to provide power to the daughterboard, we used jumpers to connect the MMB0's J5 connector to the daughterboard's J4 connector.

Finally, we glued 3/8" nylon hex standoffs to the rubber feet on the bottom of the MMB0 motherboard and used these to install it into the back of the enclosure so the DC-IN jack lined up with the access hole.

*ADS1299 daughterboard.* The daughterboard is installed in the enclosure using 7/8" hex standoffs, oriented so the J6 connector for the electrodes is close to the enclosure's face plate.

To connect the daughterboard to the enclosure's front panel: We used J6 pin 4 as the common ground and connected it to the enclosure's front panel ground banana jack. The rest of J6 comprises groups of 4 pins for each of the 8 channels. The even pins (right side) are the positive and negative inputs for each channel, which we wired to the banana jacks on the enclosure.

To connect the daughterboard to the interface PCB, we used a ribbon cable to connect the daughterboard's J3 connector to the blue 'extra' header on the interface PCB.

*Interface PCB.* The interface PCB (***Supplemental Fig 3d***, ***Supplemental Fig 4***) provided power and data to the Teensy and to an optional serial-to-USB debugger. This interface PCB was repurposed from a previous project, so many parts were not populated because they were not relevant. We are presently designing a simpler version with irrelevant components removed.

The power cable from the MMB0 connected to the X1 connector on the interface PCB, which connected to the step-down voltage regulator in IC9. To denoise the power circuitry, we placed a 220µF capacitor in C1, a 1 µF capacitor in C6, 0.1 µF capacitors in C8 and C10, and 10 µH inductors in L2 and L3. We recommend an additional 100 µF capacitor in C3 but it was not present for our recordings.

The right side of the interface PCB houses the Teensy and a 2x10 'extra' connector. The 'extra' connector is connected to the daughterboard's J3 header and to the power and data LED indicators. Instead of soldering the Teensy directly to the board, we soldered female headers for it to plug into. We used wire wrap to connect the pins of the 'extra' connector to the inputs on the Teensy (***Supplemental Fig 3e***) as described in ***Supplemental Table 1***.

Pins 2 and 4 on the daughterboard's J3 header are unused, and so we used those pins on the 'extra' header to control the power and data LEDs. To do this, we cut the unused wires of the ribbon cable and instead connected those pins on the interface PCB to the positive legs of the LEDs.

*USB debugger.* We installed a serial-to-USB debugger (Acroname S27-USB-Serial) in the grid of extra through-holes in the lower left of the interface PCB. It was powered by the 3.3V output of the Teensy and its TX and RX pins connected to Teensy pins 0 and 1 respectively. This device is optional.

**Description of the Teensy software**

The Teensy 4.0 is a 32-bit Arduino-compatible microprocessor board running at 600MHz. It was programmed using the C++ language in the Arduino IDE. The Teensy controls the ADS1299 and acts as an intermediary between the ADS1299 and a computer program, accepting commands from the computer and facilitating high speed data transfer for display, analysis and logging. We have written such programs in National Instruments' Labview and in Python for a Linux PC. The Teensy's code and Linux-based Python GUI are publicly available on our github page (https://github.com/Rodgers-PAC-Lab/ABR2025). There are several ways to upload the program to the Teensy, including from the Arduino IDE or from VSCode using the PlatformIO plugin.

The Teensy interfaces with the ADS1299 via a standard 4-wire SPI bus including chip-select (CS), serial clock (SCLK), Teensy data out (MOSI) and Teensy data in (MISO). Other ADS1299 pins required for the interface are START, RESET and data ready (DRDY).

The ADS1299 power-up sequence requires setting chip-select active (CS) then activating RESET for at least 9 µs before releasing CS. Before starting a recording, you can change the ADS1299 control registers to set the desired sampling rate and gain. Sampling rate will be the same for all channels, but each channel can have its own gain from the allowed values (1, 2, 4, 6, 8, 12, 24). We use a gain of 24 for neural channels and a gain of 1 for the audio channel.

A sampling session begins by sending the read data continuous (RDATAC) and START commands. The falling edge of DRDY signals that the latest sample is ready for reading and triggers an interrupt routine to store the new data. For example, at the highest sampling rate of 16KHz, DRDY falls every 62.5 µs. Teensy software utilizes an interrupt-driven double-buffered sampling method. All 8 channels are read at once and stored to one of the two buffers. When one buffer is full, the other buffer becomes the active buffer and the full buffer is sent over USB to the computer program. This allows data to be captured and processed continuously without dropping samples.

One 8-channel sample from the ADS1299 includes 3 status bytes plus 3 data bytes/channel, for a total of 27 bytes. Our buffers, or packets, hold 500 samples. Before sending a packet, we prepend a 12-byte header and promote samples from 24 bits to 32-bit integers, making the packet size 16,012 bytes. At 16KHz sample rate a buffer fills in 31.25 ms, for a data rate of 512 Kbytes/sec. The Teensy's USB 2.0 full speed interface has a typical maximum data rate of 30-40 Mbytes/sec, so we are well within the USB bus capability.

The packet header contains fields for packet sequence number, number of samples and channels, sampling rate, and an input trigger marker. The sequence number allows the receiving program to detect lost packets. The trigger marker is a number between 0 and 499 that marks the packet sample on which an external trigger was detected, allowing the receiving program to calculate the sample at which the trigger occurred. Currently this trigger is unused.

ADS1299 provides auxiliary functions for measuring electrode impedance and lead-off detection. By setting the appropriate control registers, an internal current source can be routed to the electrodes. Electrode impedance can be determined from the resulting voltage at the electrode. Similarly, an out of range electrode impedance can be used to detect a lead-off condition.

Transfer of data from the Teensy to the high-level program has been explained above. It uses a binary format due to the high data rate required. On the other hand, commands from the high-level program to Teensy involve a single letter to denote the command and a few numeric parameters. Simple ASCII strings suffice for this and simplify decoding on the Teensy end. Commands supported are:

U - Set up sampling. Parameters: number of channels, sample rate, flags and gain of each channel

S - Start sampling, with parameters previously given by U

I - Inquire if receiver is present. Used at program startup to make sure the Teensy and ADS1299 are operational.

Q - Query specifications. Returns the capabilities of the ADS1299, such as max sample rate, max number of channels, etc.

X - Stop sampling

After sending the Start command, the high-level program polls the USB bus to see if data is available. If so, it reads the packet header and then the raw data. The Stop command is sent when the session completes.

#### Supplemental Tables

**Supplemental Table 1. Pinout of ADS1299 daughterboard's J3 header, 'extra' cable into interface PCB, and Teensy GPIO pins.**

| ADS daughterboard J3 header OUT (2x10 header) | |
|---|---|
| **J3 pin number** | **Description** |
| 1 | **START** |
| 2 | N/C |
| **3** | **CLK** |
| 4 | N/C |
| 5 | N/C |
| 6 | N/C |
| **7** | **CS** |
| **8** | **RESET** |
| 9 | N/C |
| **10** | **GND** |
| **11** | **DIN** |
| 12 | N/C |
| **13** | **DOUT** |
| 14 | N/C |
| **15** | **DRDY** |
| 16 | N/C |
| 17 | N/C |
| 18 | N/C |
| 19 | N/C |
| 20 | N/C |

| 'Extra' cable IN to interface PCB (2x8 header) and Teensy | | |
|---|---|---|
| **'Extra' pin number** | **Description** | **Teensy pin number** |
| 1 | N/C | N/C |
| **2** | **DATA LED** | **29** |
| 3 | N/C | N/C |
| **4** | **PWR LED** | **31** |
| 5 | N/C | N/C |
| 6 | N/C | N/C |
| 7 | N/C | N/C |
| 8 | **GND** | **GND** |
| 9 | **CLK** | **13** |
| 10 | MOSI | 11 |
| 11 | MISO | 12 |
| 12 | **CS** | **33** |
| 13 | **RESET** | **24** |
| 14 | **START** | **25** |
| 15 | **DRDY** | **26** |
| 16 | N/C | N/C |

| **Other Teensy pins** | **Description** |
|---|---|
| 0 | RX1 (to serial debug's TX) |
| 1 | TX1 (to serial debug's RX) |
| 3.3V | To serial debug's VCC |
| VIN | From voltage regulator |

The top left table lists the pins in the J3 header on the ADS daughterboard. The top right table lists the pins for the 'extra' header on the interface PCB. The color coding indicates which pins are connected to each other via the ribbon cable. The bottom table indicates which other pins on the Teensy need to be connected to the serial debugger (top three rows) or motherboard power (bottom row).

### Supplemental Figures

#### Supplemental Figure 1. Hardware enclosure.

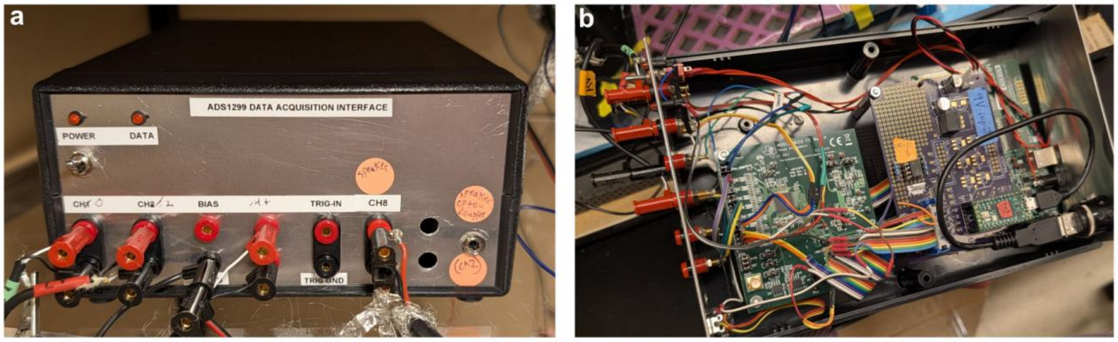


***a)*** Front panel of the enclosure.

***b)*** Inside of the enclosure with the top removed. Daughterboard (green) on left side and interface PCB (purple) above motherboard (green) on right side.

**Supplemental Figure 2. Overall schematic of the hardware inside the enclosure.**

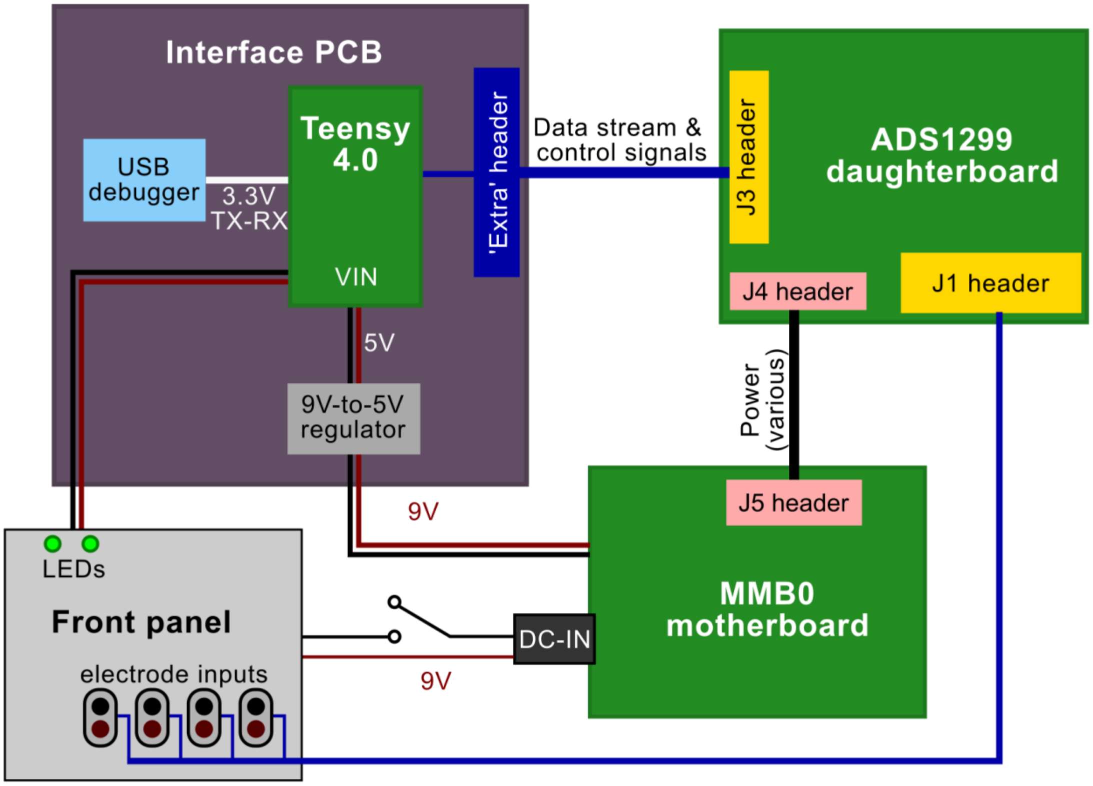


The interface PCB (purple; upper left) housed the Teensy, the USB debugger, a power regulator, and a header labeled 'extra'. The ADS1299 daughterboard (green; upper right) housed the ADS1299 itself. The MMB0 motherboard (green; lower right) provided power. The front panel (gray; lower left) provided banana jacks for signal inputs and status LEDs.

**Supplemental Figure 3. Changes to the stock hardware.**

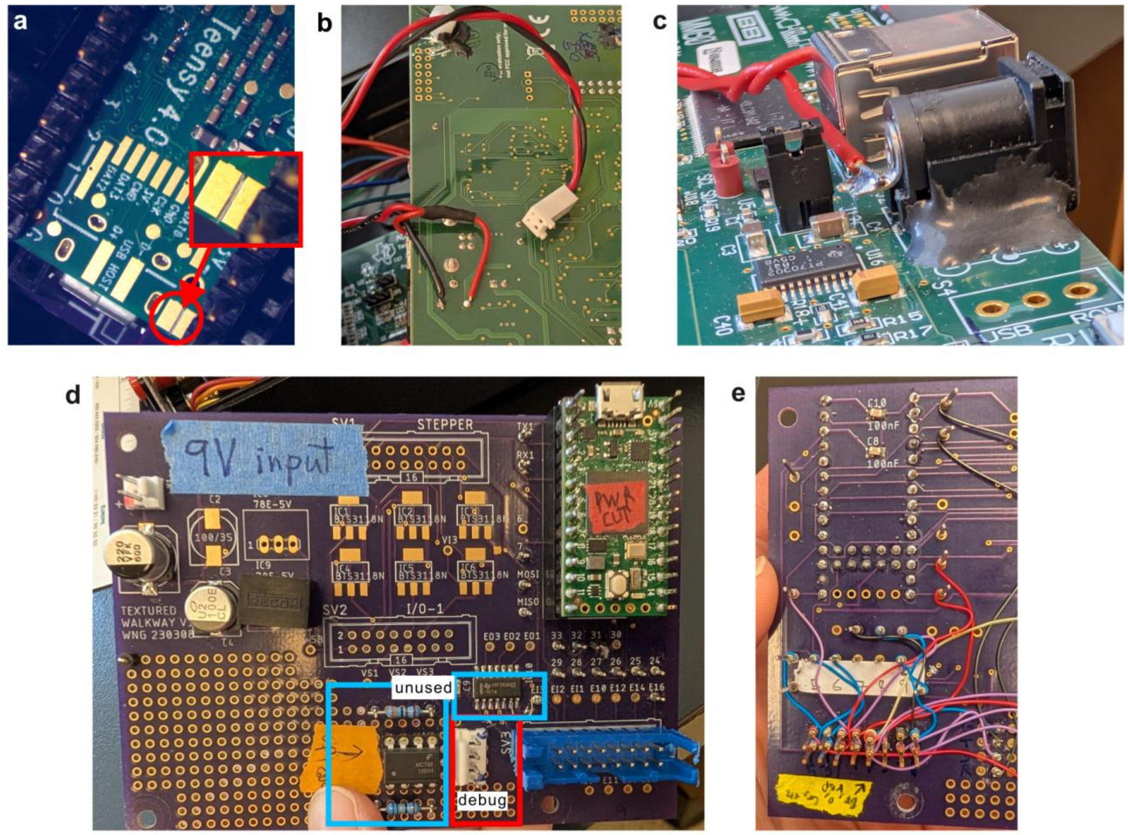


***a)*** The cut trace between the VIN and VUSB pads on the back of the board.

***b)*** Power cable to the interface PCB.

***c)*** The positive lead of the DC-IN jack is cut to allow the power switch to control power. Bending the leg away from the jack prevents it from making intermittent contact.

***d)*** Front of the interface PCB. The header for the optional serial debugger is outlined in red. The areas outlined in blue were populated but were not used in the final version of the device and so they are not described in this manuscript.

***e)*** Back of the interface PCB. Wire wrap connected ‘extra’ header pins to Teensy pins as shown in ***Supplemental Table 1.***

**Supplemental Figure 4. Layout of interface PCB.**

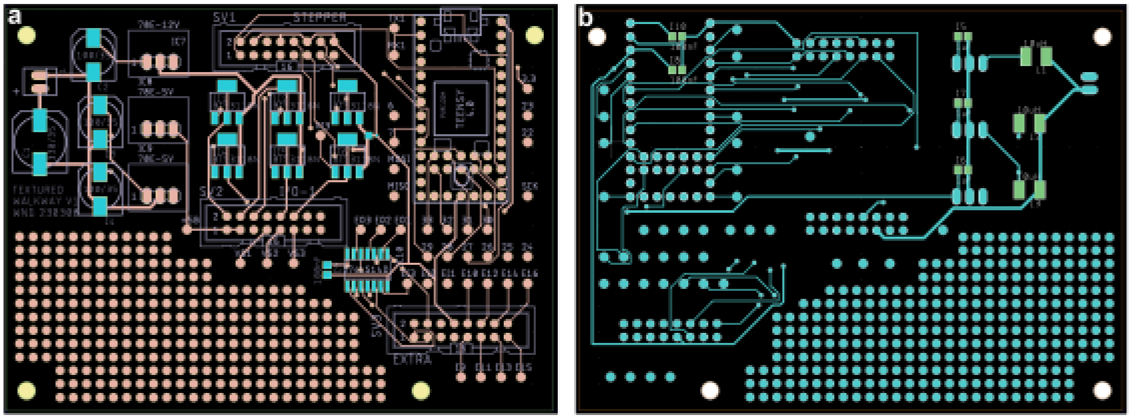


***a)*** Layout of the front of the interface PCB.

***b)*** Layout of the back of the interface PCB. The design files for this PCB are available on our github.
